# Inhibition of glycosphingolipid biosynthesis reverts multidrug resistance by differential modulation of ABC transporters on chronic myeloid leukemias

**DOI:** 10.1101/2020.02.18.954297

**Authors:** Eduardo J. Salustiano, Kelli M. da Costa, Leonardo Freire-de-Lima, Lucia Mendonça-Previato, José O. Previato

## Abstract

Multidrug resistance (MDR) in cancer manifests due to cross-resistance to chemotherapeutic drugs with neither structural nor functional relationship, markedly by increased expression and activity of ABC superfamily transporters. Evidences indicate sphingolipids as substrates to ABC proteins in processes such as cell signaling, membrane biosynthesis and inflammation, and products of its biosynthetic route were shown to favor cancer progression. Glucosylceramide (GlcCer) is a ubiquitous glycosphingolipid (GSL) generated by glucosylceramide synthase, a key cell regulator enzyme encoded by the UDP-glucose ceramide glucosyltransferase (UGCG) gene. Under stress, cells increase *de novo* biosynthesis of ceramides, which return to sub-toxic levels after assimilation into GlcCer by UGCG. Given that cancer cells seem to mobilize UGCG and increase GSL contents for the clearance of ceramides ultimately contributing to treatment failure, we studied how inhibiting GSL biosynthesis would affect the MDR phenotype of chronic myeloid leukemias. Results indicate that MDR associates to higher expression of UGCG and to a complex GSL profile. Inhibition of this glucosyltransferase greatly reduced GM1 expression, and cotreatment with standard chemotherapeutics sensitized cells leading to mitochondrial membrane potential loss and apoptosis. Despite reducing ABCB1 expression, only the ABCC-mediated efflux activity was affected. Consistently, efflux of C6-ceramide, one byproduct of UGCG downregulation, was reduced after inhibition of ABCC-mediated transport. Overall, UGCG inhibition impaired the malignant glycophenotype of MDR leukemias, overcoming drug resistance through distinct mechanisms. This work brings more comprehension about the involvement of GSL for chemotherapy failure, and modulation of its contents emerges as an intervention targeted to MDR leukemias.

---

As a multifactorial sum of diseases, cancer presents great challenges to the development of safe, successful therapies. Distinct mechanisms employed by transformed cells to avoid toxicity generated by chemotherapy often crosstalk, leading to an adapted phenotype comprising both intrinsic and acquired drug resistance. Multidrug resistance (MDR) is the main hurdle to chemotherapy success, as stress-adapted molecular mechanisms including reduced influx, increased efflux and accelerated metabolism of xenobiotics work in tandem reducing the effective concentration at the molecular target (1). Efflux transporters such as ATP-binding cassette (ABC) proteins actively detoxify cells and tissues from both xenobiotics and toxic metabolites, playing major roles in MDR. Regardless of the diversity of ABC subfamilies and isoforms two proteins are mostly associated to MDR, ABCB1 (P-glycoprotein, Pgp) and ABCC1 (Multidrug Resistance Protein 1, MRP1) (1, 2). ABCB1 and ABCC1 share similarities and differences when cellular localization and substrate specificity are considered. The latter is mostly located on plasma membrane, the first is present on any cell membrane, and both were shown to associate to microdomains such as lipid rafts depending on cell subtype (3, 4). Both actively extrude a variety of non-related chemotherapeutic drugs, but ABCC1 is able to transport substrates in conjugation with glutathione (GSH) or in cotransport as well (5), playing complementary roles for the MDR phenotype. ABCB1 interacts with liposoluble or amphipathic molecules that are prone to accumulate in the intramembrane space (6), and ABCC1 exhibit higher affinity for negatively-charged glucoronates, sulfates, GSH-conjugated compounds and products from lipid metabolism (7).

Sphingolipids such as ceramides and its phosphorylated or glycosylated forms are directly involved in cell fate, assuming active parts on either cell proliferation or death (8). Ceramides, whether originated from sphingomyelin remodeling or synthesized *de novo* on the endoplasmic reticulum (ER) are transferred to cis-Golgi, where they are employed as substrates to UDP-glucose ceramide glucosyltransferase (UGCG) to form glucosylceramide (GlcCer), the precursor to all glycosphingolipids (GSL). Endogenous ceramides have been directly linked to cancer treatment, given that chemotherapeutic agents with unrelated mechanisms for example paclitaxel, daunorubicin, etoposide (9-11) and the tyrosine kinase inhibitors sorafenib and imatinib (12) rely on ceramides to drive the intrinsic pathway of apoptosis through caspase activation or caspase- and p53-independent mitotic catastrophe (11, 13). Second to their structural role on the organization of lipid rafts (14), GSL relates to development of drug resistance considering that cancer cells often present increased UGCG expression, being able to incorporate ceramides on GSL (15). Concerning MDR, a close crosstalk of ABCB1 and GSL has been observed; ABCB1 and UGCG were coincidently overexpressed in drug-resistant breast, ovary, cervical, colon cancer and on chronic myeloid leukemias (16, 17); GlcCer regulates ABCB1 expression through Wnt/β-catenin and cSrc signaling (18); and this transporter is able to act as a flippase on the transfer of GlcCer from the cis-Golgi to trans-Golgi during GSL biosynthesis (18). Despite its capacity of translocating sphingolipids such as sphingosine-1-phosphate (19) and GlcCer on polarized cells (20) and its coexpression with UGCG on colon cancer (21), similar relationship involving ABCC1 activity and GSL is not clear.

Considering the diversity of mechanisms MDR cancer cells resort to avoid and adapt to chemotherapeutic stress and the prime involvement of UGCG on the generation of GSL (22), the fate of endogenous ceramides is critical to successful cancer chemotherapy on a molecular level. Several studies evaluated the expression of ABCB1 and reversal of drug sensitivity on solid tumors and its association with GSL; nevertheless, our work focused on leukemic cells that express both ABCB1 and ABCC1, extending to the functional evaluation of those proteins after UGCG inhibition, which finds little coverage from the literature. In this context, we report the distinct ways ABCB1 and ABCC1 expression and activity were modulated after impairment of GSL biosynthesis on clinically relevant models of drug-resistant chronic myeloid leukemias.

## Results

### MDR chronic myeloid leukemias overexpress UGCG along with a complex GSL profile, which is reverted after treatment with a ceramide analogue

*De novo* ceramide synthesis on Golgi increases during stress, and cancer cells are able to upregulate ceramide glycosylation ultimately changing GSL contents on cell membranes. To determine if selection with standard chemotherapeutics would alter these processes on human leukemias, the expression of UGCG, and profiles of GSL and GM1 were evaluated on K562 (drug-sensitive), and on MDR derivatives Lucena-1 (K562/VCR) and FEPS (K562/DNR) cells. Results on Fig. 1A and 1B indicate that the glucosyltransferase that catalyzes the addition of UDP-glucose to ceramide, UGCG, is upregulated on the MDR leukemia models, notably on the FEPS line. Next, the three cell lines were treated with a specific UGCG inhibitor, the ceramide analogue EtDO-P4, and their effects on viability and GSL contents are depicted on Fig. 1C, 1D and 1E. Based on these results, a sub-toxic concentration of this inhibitor was employed for further assays. MDR cells showed increased contents and more complex profiles of GSL after thin-layer chromatography when compared to the parental, and treatment with 1 μM EtDO-P4 for 24 h significantly reduced GSL expression on all three leukemias. Since total extraction does not distinguish the GSL on plasma or intracellular membranes, log-phase growing, viable cells were stained with cholera toxin (CHT-FITC). CHT specifically binds GM1 on the extracellular leaflet, and an increase in its fluorescence could be observed on the MDR cells Lucena-1 and FEPS after flow cytometry. In accordance, UGCG inhibition reduced GM1 contents in 55% and 75% on these cells, contrasting to a reduction of 35% on drug-sensitive K562 (Fig. 1F and 1G).

**Figure 1.**
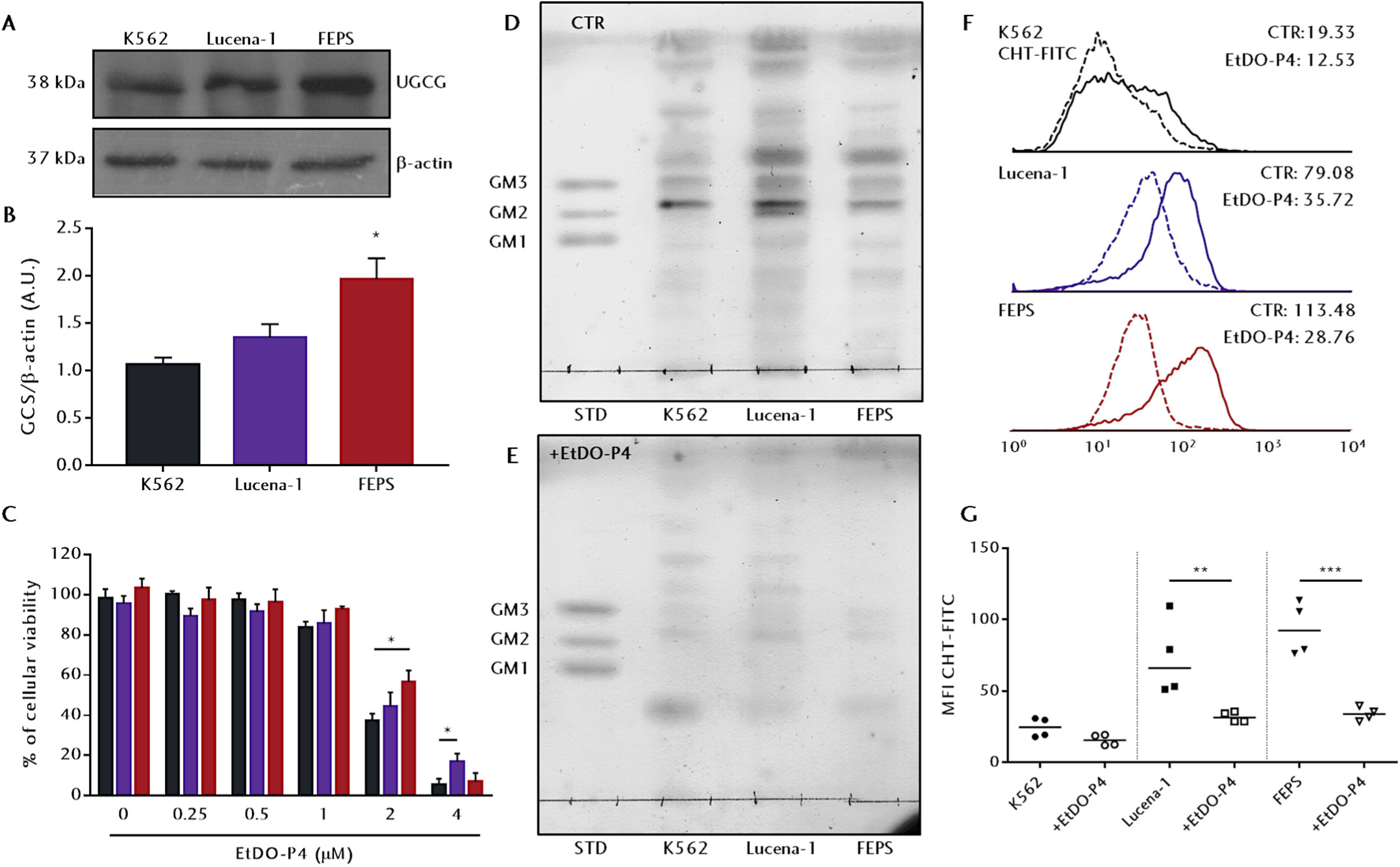
UDP-glucose ceramide glucosyltransferase (UGCG) and glycosphingolipid expression on human erythroleukemias, and effect of UGCG inhibition on those profiles. K562 and their MDR counterparts Lucena-1 and FEPS were cultured at concentrations of 2×10^4^ cells/mL for 72 h in presence or absence of EtDO-P4. (**A**) UGCG expression was analyzed by Western blotting as described in Experimental Procedures. Representative images from four independent extractions. (**B**) Band densities were quantified, and the amount of UGCG was calculated as the density of the UGCG band divided by the density of the β-actin band for each cell line. Bars represent the mean UGCG to β-actin ratios + SD. n=4. (**C**) Black, violet or red bars respectively represent the mean percentages of cellular viabilities + SEM for K562, Lucena-1 or FEPS, as measured by MTT assay. Normalized data from three independent experiments, with each concentration evaluated in triplicate. (**D**) and (**E**) Total lipids from 5×10^7^ cells were extracted, purified, resolved by TLC and developed by resorcinol/HCl staining as indicated under Experimental Procedures. The migration pattern of a GSL standard containing a mix of GM1, GM2 and GM3 is indicated on the left of the image (STD). Controls (CTR) were treated with diluent. Representative images from two independent extractions. (**F**) K562, Lucena-1 and FEPS were incubated for 1 h with 5 μg.mL^-1^ FITC-cholera toxin (CHT-FITC), and the presence of the prototype monosialotetrahexosylganglioside GM1 was assessed by the median fluorescence intensity (MFI) of viable cells in a flow cytometer. Representative histograms for the MFI of whole populations of treated (dashed lines) and untreated populations (continuous lines). (**G**) Scatter plots for the MFI obtained from each cell line as follows: black circles for K562, squares for Lucena-1 and inverted triangles for FEPS untreated cells, and white ones for EtDO-P4 treatment. n=4. **\*p<0.05**, \***\*p<0.01** and **\*\*\*p<0.001**.

### Pharmacological UGCG inhibition induces cytotoxicity and decreases drug resistance

Disruption of the GSL biosynthesis machinery could lead to accumulation of sphingolipid mediators, altering cell signaling and leading to either survival or death depending on the intrinsic properties of each cell subtype. Thus, the effects of UGCG inhibition on cellular viability are depicted on Table 1. Results indicate that the MDR phenotype and the increased UGCG expression translated into a modest resistance to EtDO-P4, since only FEPS presented higher viability and IC_50_ after a 72 h treatment (Fig. 1C and Table 1).

**Table 1.** Cytotoxicity of standard drugs before and after UGCG inhibition. K562, Lucena-1 and FEPS cells were co-incubated for 72 h with a range of concentrations of either vincristine (VCR), daunorubicin (DNR) or cisplatin (CDDP) and 1 μM EtDO-P4, and compared to ones treated with EtDO-P4 and C6-ceramide (C6-cer). Viability was evaluated by MTT assay, and results are reported as mean concentrations giving half-maximal inhibition (IC_50_) ± SD, in micromolar (μM). Results obtained from three independent experiments, with each concentration evaluated in triplicate. The **¥** symbols denote comparisons to untreated cells whereas **†** indicate comparison to the parental cell K562. n=3, with **\*p<0.05** and **\*\*p<0.01**.

| IC<sub>50</sub> | K562 | +EtDO-P4 | Lucena-1 | +EtDO-P4 | FEPS | +EtDO-P4 |
| --- | --- | --- | --- | --- | --- | --- |
| VCR | 0.054±0.005 | 0.043±0.001 | 0.85±0.12 | 0.45±0.07*,¥ | 1.05±0.07 | 1.02±0.08 |
| DNR | 0.082±0.001 | 0.076±0.007 | 3.06±0.87 | 0.66±0.13**,¥ | 6.45±1.27 | 2.75±0.42**,¥ |
| CDDP | 2.83±0.22 | 3.97±0.50 | 6.07±0.48 | 4.99±0.09 | 7.41±0.62 | 5.07±0.48*,¥ |
| EtDO-P4 | 1.87±0.13 | - | 2.14±0.59 | - | 2.50±0.25*,† | - |
| C6-cer | 39.60±5.00 | - | 22.20±2.44*,† | - | 18.73±4.70*,† | - |

Given that UGCG is the first enzyme on the GSL biosynthesis pathway, inhibition of its activity would impair cell responses to stress. In this context, results on Table 1 suggest that GSL depletion synergizes with the cytotoxicity caused by chemotherapeutic drugs with diverse mechanisms of action, as sub-lethal cotreatment with EtDO-P4 reduced the IC_50_ for vincristine (VCR) and daunorubicin (DNR) on Lucena-1 and the IC_50_ for DNR and cisplatin (CDDP) on FEPS cells. This reduction did not manifest on K562, which in line with earlier results, suggests that GSL depletion showed minimal effects on sensitive cells. Noteworthy, treatment with one subproduct from UGCG inhibition, N-hexanoyl-D-erythro-sphingosine (C6-ceramide, C6-cer), induced opposite effect to EtDO-P4 on these cells. The lower IC_50_ values for C6-cer on Lucena-1 and FEPS than on K562 suggest that this sphingolipid induces collateral sensitivity, a hypersensitivity towards secondary agents that arises from the development of resistance towards an unrelated primary drug.

### UGCG inhibition reduces mitochondrial membrane potential (Δψ_m_) and induces apoptotic cell death

Among sphingolipids, ceramides are linked to apoptotic cell death through the intrinsic pathway due to direct or indirect interaction with the mitochondrial permeability transition pore. To investigate if the mitochondria would be involved on the cell death observed after GSL depletion, the MDR leukemias Lucena-1 and FEPS were treated with EtDO-P4, DNR or C6-cer for 24 h and incubated with rhodamine 123 (Rho123). This fluorescent dye accumulates within energized mitochondria, and this retention is progressively lost as Δψ_m_ is reduced. Fig. 2A and 2B indicate that only 2 μM EtDO-P4 significantly reduced mitochondrial Rho123 fluorescence, a similar result obtained after C6-cer treatment. As expected, DNR was not able to change mitochondria polarization; however, when combined with 1 μM EtDO-P4, Δψ_m_ was reduced to similar levels to after 2 μM EtDO-P4 or 20 μM C6-cer treatment. On par with previous results, the stress after UGCG inhibition led MDR cells to apoptosis in a dose-dependent fashion, given that 57.09% and 34.06% of Lucena-1 and FEPS, respectively, underwent early or late apoptosis (upper right + lower right quadrants) after 72 h when treated with 2 μM EtDO-P4, and over 90% after 4 μM (Fig. 2C and 2D).

**Figure 2.**
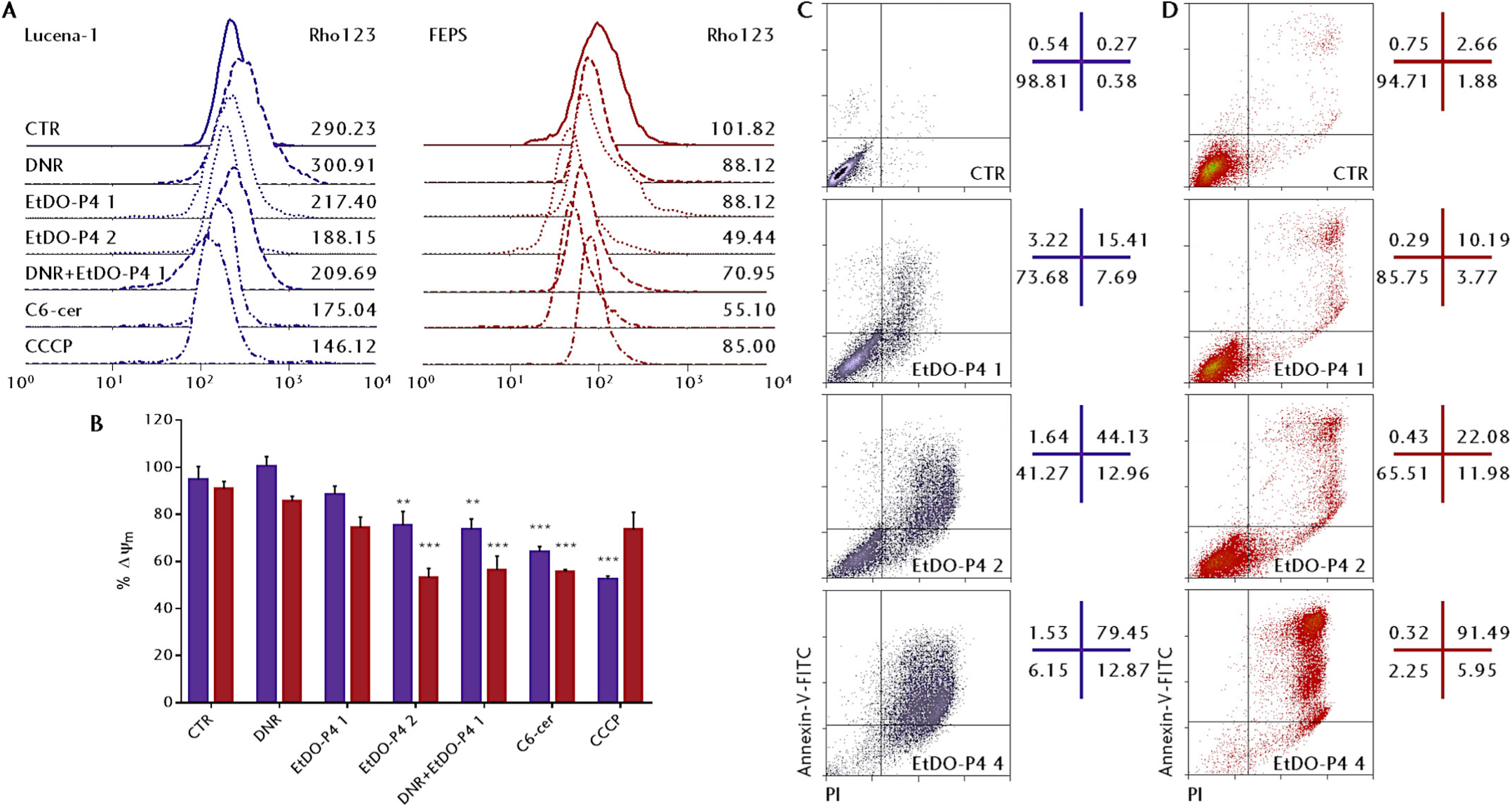
Changes in mitochondrial membrane potential (Δψ_m_) and apoptotic cell death after UGCG inhibition. The MDR cells Lucena-1 and FEPS were treated with 1 μM EtDO-P4, 1 μM daunorubicin (DNR), a combination of the two, 2 μM EtDO-P4 or with 20 μM C6-cer for 24 h. Cells were then incubated with 2.5 μM rhodamine 123 (Rho123) for 30 min at 37 °C and Δψ_m_ was estimated by the MFI of viable cells normalized to the one for untreated cells. Controls (CTR) were treated with diluent, and positive controls were treated with 50 mM CCCP for 30 min prior to Rho123. (**A**) Representative histograms for Rho123 fluorescence after each indicated treatment. Violet or red lines represent, respectively, Lucena-1 or FEPS treatments as follows: continuous lines for Rho123 (CTR); dashed lines for DNR-treated cells; dotted for EtDO-P4 treatment; and dash-dotted for the C6-cer and CCCP treatments. Numbers indicate the MFI for each condition. (**B**) Violet or red bars represent the mean MFI percentages + SEM of Lucena-1 and FEPS normalized to untreated controls. n=5, with \***\*p<0.01** and **\*\*\*p<0.001**. (**C**) and (**D**) MDR cells were cultured with diluent or 1, 2 or 4 μM EtDO-P4 for 72 h, and cell death was evaluated by Annexin V-FITC/iodidium propide (PI) double staining. Dot-plots were divided into four quadrants as follows: upper left (PI+/Annexin-V-), necrotic cells; upper right (PI+/Annexin-V+), late apoptotic cells; lower left (PI-/Annexin-V-), viable cells; lower right (PI-/Annexin-V+), early apoptotic cells. Representative dot-plots from three independent experiments.

### Differential contribution of ABC transporters to the reversal of the MDR phenotype after UGCG inhibition

Results so far showed that depletion of GSL led to increased cell death, and this synergizes with chemotherapeutic drugs likely due to accumulation of ceramides on MDR leukemias. ABCB1 and ABCC1 transporters are key drivers to MDR phenotype, actively extruding xenobiotics thus reducing cell death. In this context, the expression of these proteins was evaluated in conditions matching previous assays. Treatment with EtDO-P4 produced, as demonstrated on Fig. 3, distinct effects on expression of ABCB1 and ABCC1. Sub-toxic UGCG inhibition altered the expression of neither ABC protein, whereas treatment with a 2 μM concentration of EtDO-P4 for 24 h reduced only ABCB1 expression on both cell lines. It should be pointed that despite 2 μM EtDO-P4 induced cell stress on diverse assays after 72 h, cells were still viable after 24 h (Fig. S1).

**Figure 3.**
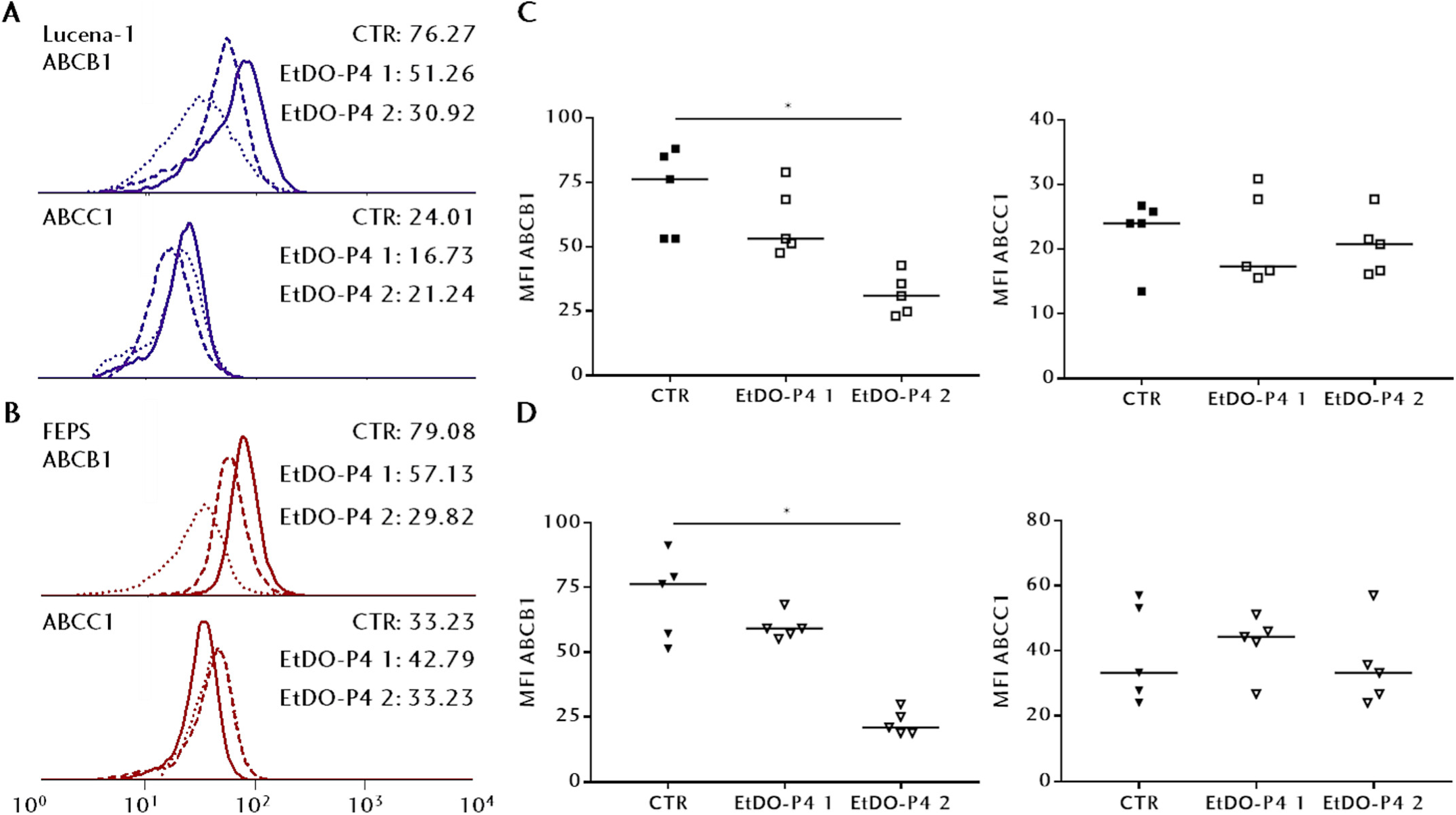
Expression of ABC MDR proteins after UGCG inhibition. MDR cells were treated with EtDO-P4 for 24 h prior to incubation with specific anti-ABCB1 and anti-ABCC1 antibodies. (**A**) and (**B**) Continuous lines represent untreated cells (CTR), whereas dashed or dotted lines represent, respectively, cells treated with 1 μM or 2 μM EtDO-P4. Controls (CTR) were treated with diluent. The median fluorescence intensity (MFI) for whole populations are present on the right. (**C**) Scatter plots for the MFI obtained from Lucena-1 or (**D**) for FEPS for ABCB1 and ABCC1, as follows: black squares or inverted triangles, respectively, for Lucena-1 and FEPS untreated cells, and white ones for EtDO-P4 treatment. Lines indicate the median ABCB1 or ABCC1 expression. n=5, with **\*p<0.05**.

Regardless of not altering ABCB1 or ABCC1 expression on mild conditions, the ABC-mediated transport after pre-treatment with EtDO-P4 was investigated as well. For this, cells were then incubated with specific fluorescent substrates for ABCB1 and ABCC transporters, and dye retention after free or inhibited efflux were analyzed by the median fluorescence intensity (MFI). Again, results in Fig. 4 indicate that GSL depletion did not impair ABCB1-mediated transport, since profiles of Rho123 MFI were similar irrespective of EtDO-P4 treatment. ABCC activity, on the other hand, was significantly modulated after treatment, since carboxyfluorescein (CF) MFI was higher after both free and inhibited efflux for Lucena-1 (Fig. 5A and 5C) and for FEPS (Fig. 5B and 5D). It is important to note that CF may be transported by other ABCC subfamily members than ABCC1, but not by ABCB1, given that ABCB1 inhibition with VP did not affect CF MFI, despite the MDR leukemias expressing both ABCB1 and ABCC1 (Fig. S2).

**Figure 4.**
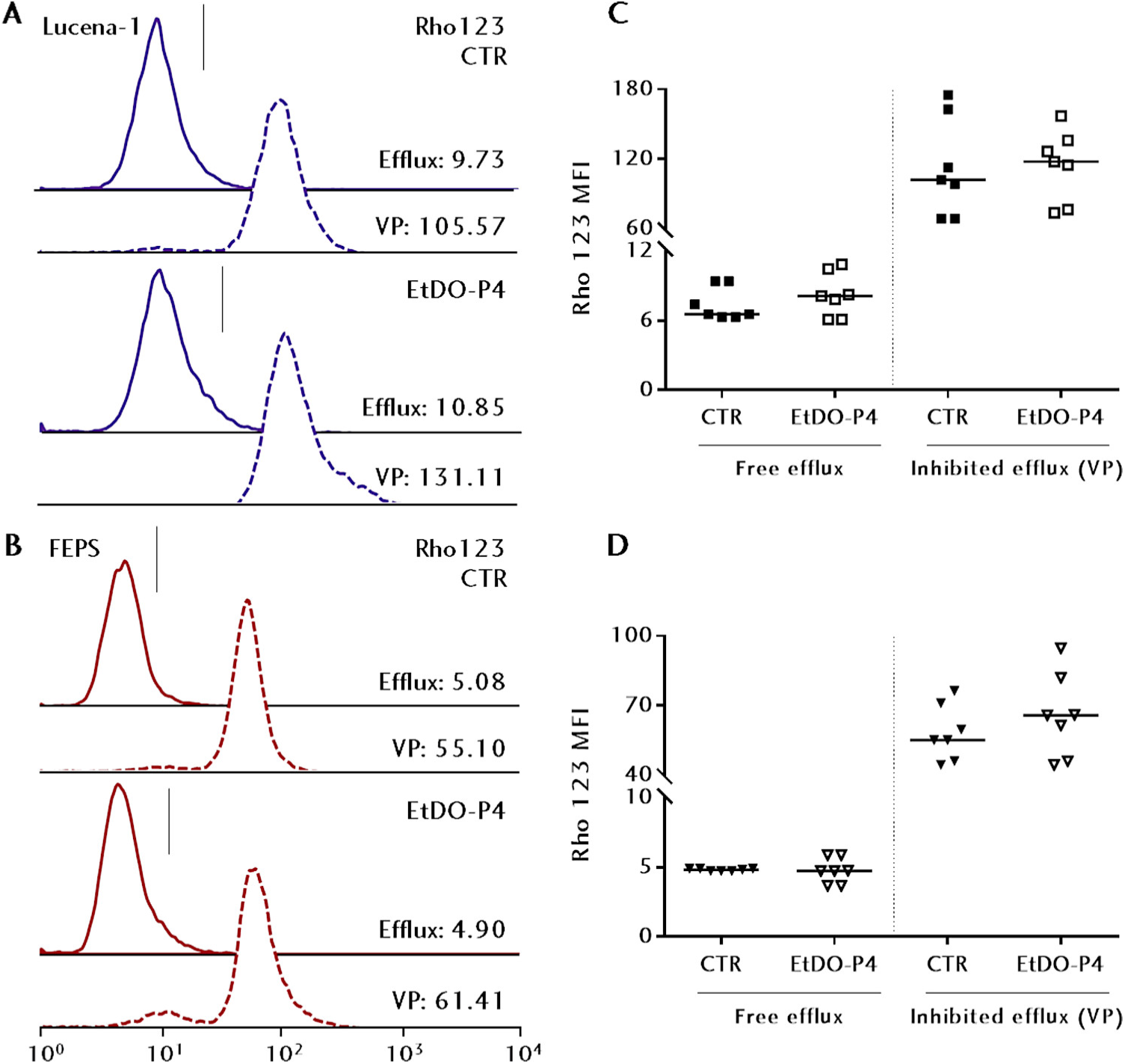
Profiles of ABCB1-mediated transport after UGCG inhibition. The transport activity by ABCB1 was evaluated by rhodamine 123 (Rho123) efflux assays. MDR cells were treated with 1 μM EtDO-P4 for 24 h, then incubated in medium containing 250 nM Rho123 for 30 min. Fresh media was added in the absence or presence of 10 μM verapamil (VP), inhibitor for ABCB1-mediated transport, for another 30 min. After incubation, the median fluorescence intensity (MFI) accounting for intracellular Rho123 was acquired by flow cytometry. Representative histograms for (**A**) Lucena-1 or (**B**) FEPS were divided into two areas separated by the vertical lines: Rho123-negative on the left and Rho123-positive on the right, as described under Experimental Procedures. Continuous or dashed lines indicate cells, respectively, in absence (free efflux) and in presence of VP (inhibited efflux). Controls (CTR) were treated with diluent. The MFI for whole populations are present on the right. Scatter plots for the MFI obtained for (**C**) Lucena-1 or (**D**) FEPS after the free or inhibited Rho123 efflux, with lines indicating the median of each population. Black squares or inverted triangles respectively indicate Lucena-1 or FEPS untreated cells, and white ones, EtDO-P4 treatment. n=7.

**Figure 5.**
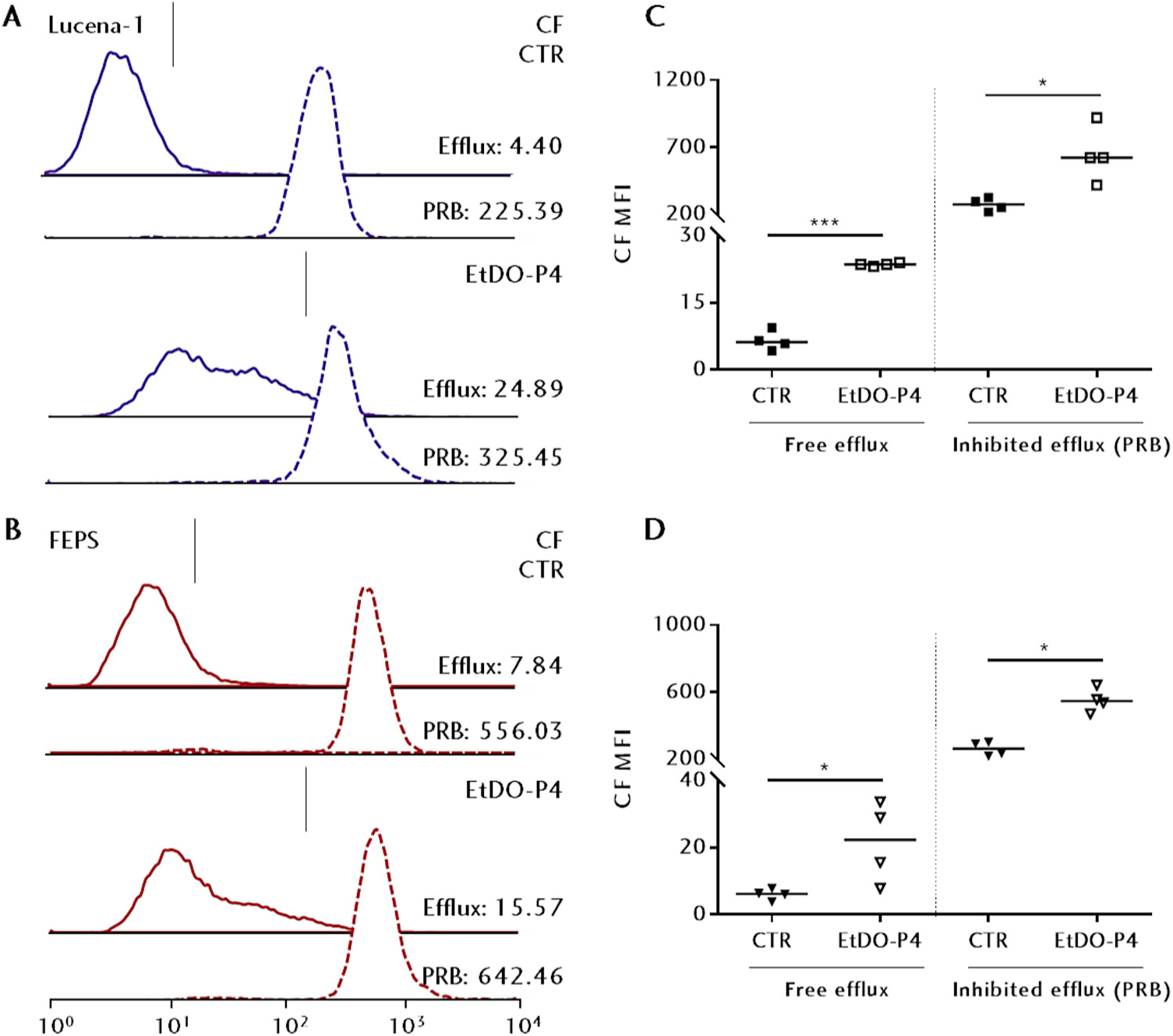
Profiles of ABCC-mediated transport after UGCG inhibition. The transport activity by ABCC subfamily members was evaluated by carboxyfluorescein (CF) efflux assays. MDR cells were treated with 1 μM EtDO-P4 for 24 h, then incubated in medium containing 500 nM CFDA for 30 min. Fresh media was added in the absence or presence of 1.25 mM probenecid (PRB), inhibitor for ABCC-mediated transport, for another 30 min. After incubation, the MFI accounting for intracellular CF was acquired by flow cytometry. Representative histograms for (**A**) Lucena-1 or (**B**) FEPS were divided into two areas separated by the vertical lines: CF-negative on the left and CF-positive on the right, as described under Experimental Procedures. Continuous or dashed lines indicate cells, respectively, in absence (free efflux) and in presence of PRB (inhibited efflux). Controls (CTR) were treated with diluent. The MFI for whole populations are present on the right. Scatter plots for the MFI obtained for (**C**) Lucena-1 or (**D**) FEPS after the free or inhibited CF efflux, with lines indicating the median of each population. Black squares or inverted triangles respectively indicate Lucena-1 or FEPS untreated cells, and white ones, EtDO-P4 treatment. n=4, with \***p<0.05** and **\*\*p<0.01**.

### ABCC but not ABCB1 transports one byproduct of a compromised GSL biosynthesis pathway

The roles of GSL and ABC transporters for the adaptation to cellular stresses are well discussed, and pharmacological impairment of UGCG was demonstrated to efficiently modulate the main features of the MDR phenotype. However, when analyzed as whole, results point to MDR leukemias being able to actively reduce ceramide levels by mechanisms apart from glycosylation. To examine this possibility, the efflux assays of ABCB1 and ABCC substrates were performed on untreated cells in the presence of C6-cer as competitive inhibitor, and the resultant MFI and percentages of cells loaded with either Rho123 or CF after the efflux phases were evaluated. In agreement with previous experiments, one more time the ABCB1 efflux of Rho123 was not altered in the presence of increasing C6-cer concentrations (Fig. 6A-D and Fig. S3A). An opposite outcome was observed when C6-cer was co-incubated during the CF efflux assay, as results indicate CF retention in both Lucena-1 and FEPS. MFI was dose-dependently increased on both MDR cells, notably when 40 μM C6-cer was employed as a competitor to ABCC-mediated efflux (Fig. 7A-D). This profile could be better appreciated when observing the percentages of CF+ cells, which reached 50% in the presence of 40 μM C6-cer (Fig. S3B). These results suggest that C6-cer could be transported by ABCC proteins.

**Figure 6.**
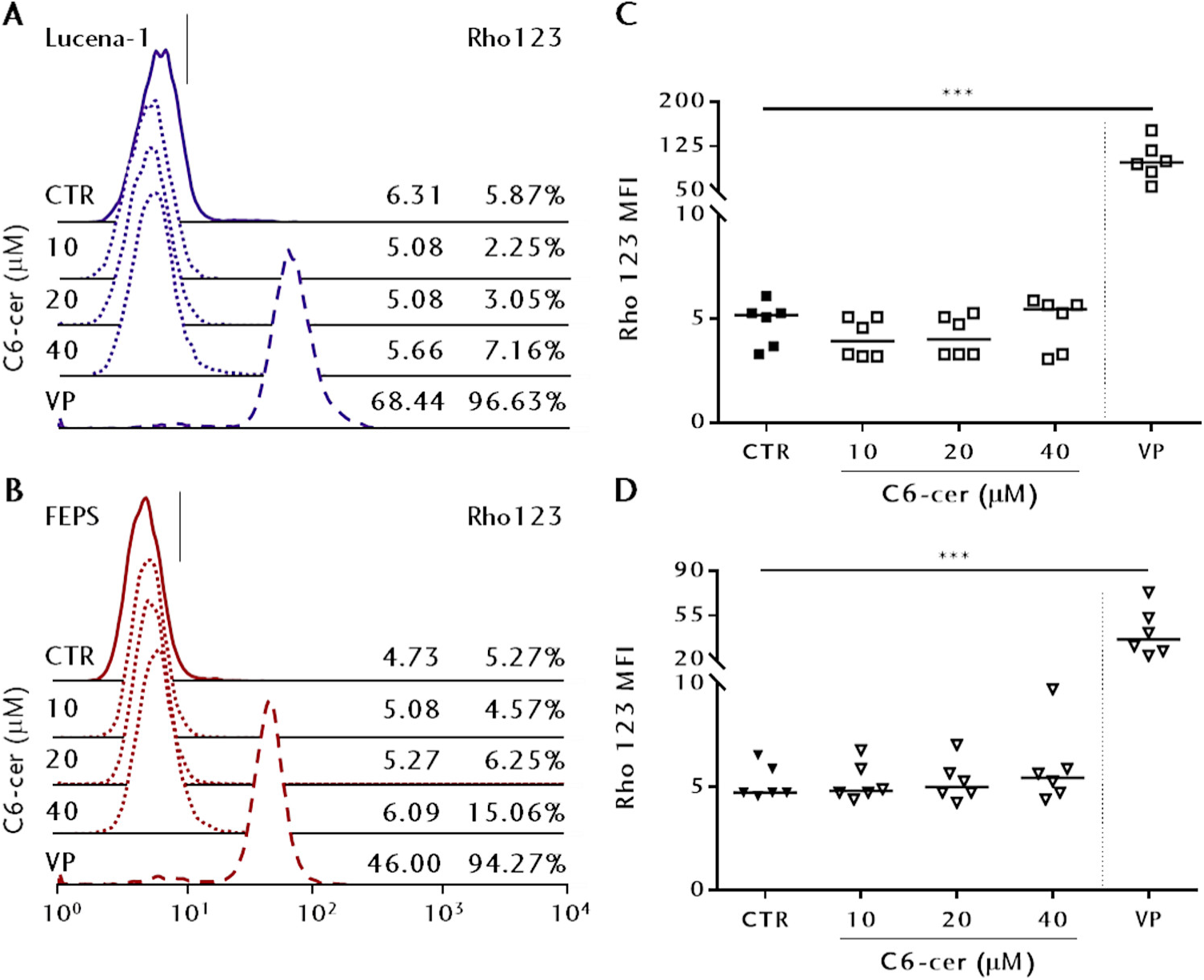
Lack of modulation of ABCB1-mediated efflux by one byproduct from a defective GSL biosynthesis pathway. The effect of C6-ceramide competition for ABCB1-mediated transport was evaluated by rhodamine 123 (Rho123) efflux assays. MDR cells were incubated in medium containing 250 nM Rho123 for 30 min. Fresh media was added in the absence or presence of 10 μM verapamil (VP), inhibitor for ABCB1-mediated transport, or of a range of concentrations of C6-cer for another 30 min. After incubation, the median fluorescence intensity (MFI) accounting for intracellular Rho123 was acquired by flow cytometry. Representative histograms for (**A**) Lucena-1 or (**B**) FEPS were divided into two areas separated by the vertical lines: Rho123-negative on the left and Rho123-positive on the right, as described under Experimental Procedures. Continuous, dotted or dashed lines indicate cells, respectively, in absence (CTR, free efflux) and in presence of C6-cer, or VP (inhibited efflux). Controls (CTR) were treated with diluent. The MFI for whole populations and percentages of Rho123-positive cells are listed on the right. Scatter plots for the MFI obtained for (**C**) Lucena-1 or (**D**) FEPS after the free, C6-cer-competitive or inhibited Rho213 efflux. Black squares or inverted triangles respectively indicate Lucena-1 or FEPS untreated cells, and white ones, C6-cer or VP treatment, with lines indicating the median of each population. n=6, with **\*\*\*p<0.001**.

**Figure 7.**
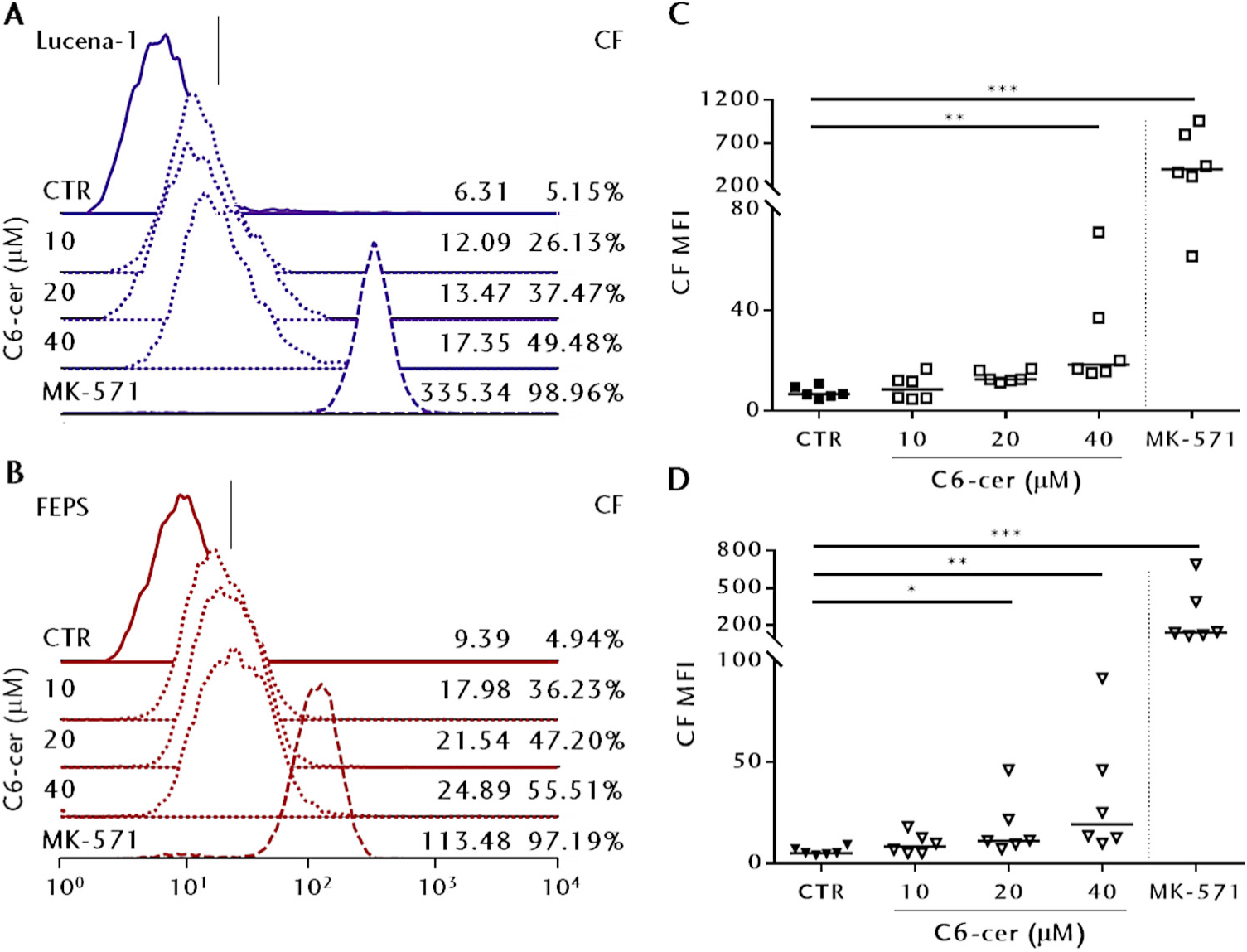
Modulation of ABCC-mediated efflux by one byproduct from a defective GSL biosynthesis pathway. The effect of C6-ceramide competition for ABCC-mediated transport was evaluated by carboxyfluorescein (CF) efflux assays. MDR cells were incubated in medium containing 500 nM CFDA for 30 min. Fresh media was added in the absence or presence of 25 μM MK-571, inhibitor for ABCB1-mediated transport, or of a range of concentrations of C6-cer for another 30 min. After incubation, the median fluorescence intensity (MFI) accounting for intracellular Rho123 was acquired by flow cytometry. Representative histograms for (**A**) Lucena-1 or (**B**) FEPS were divided into two areas separated by the vertical lines: CF-negative on the left and CF-positive on the right, as described under Experimental Procedures. Continuous, dotted or dashed lines indicate cells, respectively, in absence (CTR, free efflux) and in presence of C6-cer, or MK-571 (inhibited efflux). Controls (CTR) were treated with diluent. The MFI for whole populations and percentages of CF-positive cells are listed on the right. Scatter plots for the MFI obtained for (**C**) Lucena-1 or (**D**) FEPS after the free, C6-cer-competitive or inhibited CF efflux. Black squares or inverted triangles respectively indicate Lucena-1 or FEPS untreated cells, and white ones, C6-cer or MK-571 treatment, with lines indicating the median of each population. n=6, with **\*p<0.05**, \***\*p<0.01** and **\*\*\*p<0.001**.

To further delineate if ceramides would be directly transported out by ABC proteins, the activity assay was once more performed on untreated cells, this time with a fluorescent, nitrobenzodiaxole-labeled ceramide derivative (C6-NBD-cer) as substrate. Results on Fig. 8 corroborate that ABCC transporters mediate the efflux of C6-NBD-cer, owing to fact that the MFI and percentages of C6-NBD-cer+ cells significantly increased in the presence of MK-571 but not VP when compared to controls (Fig. 8A-D and Fig. S3C).

**Figure 8.**
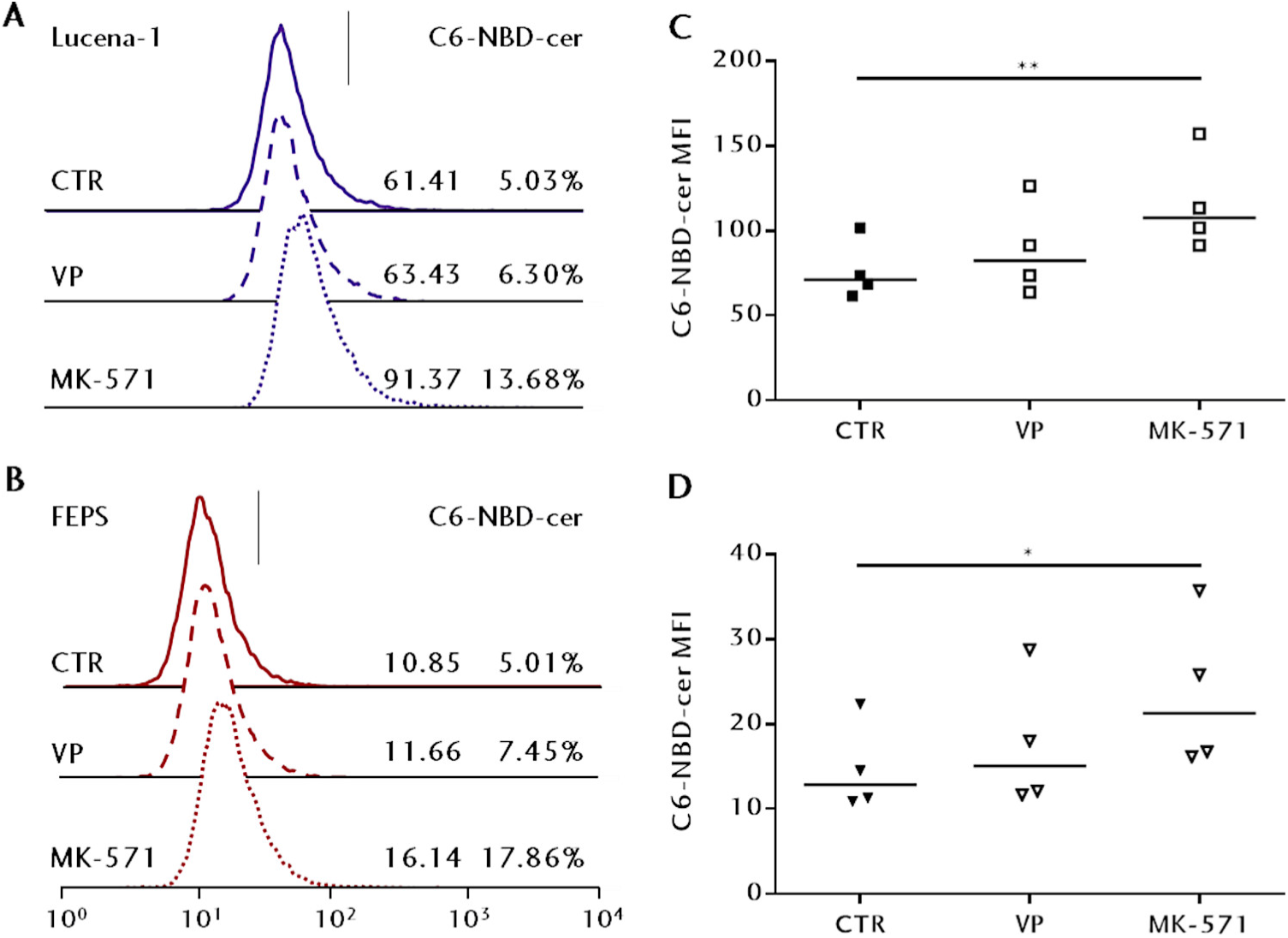
ABC-mediated efflux of a fluorescent ceramide derivative. The ABC-mediated transport was evaluated by nitrobenzoxadiazole-labeled C6-ceramide (C6-NBD-cer) efflux assays. MDR cells were incubated in medium containing 1 μM C6-NBD-cer for 30 min. Fresh media was added in the absence or presence of 10 μM VP or 25 μM MK-571, standard ABCB1 or ABCC inhibitors, for another 30 min. After incubation, the median fluorescence intensity (MFI) accounting for intracellular Rho123 was acquired by flow cytometry. Representative histograms for (**A**) Lucena-1 or (**B**) FEPS were divided into two areas separated by the vertical lines: C6-NBD-cer-negative on the left and C6-NBD-cer-positive on the right, as described under Experimental Procedures. Continuous, dashed or dotted lines indicate cells, respectively, in absence (CTR, free efflux) and in presence of VP (ABCB1 inhibited efflux) or MK-571 (ABCC inhibited efflux). Controls (CTR) were treated with diluent. The MFI for whole populations and percentages of CF-positive cells are listed on the right. Scatter plots for the MFI obtained for (**C**) Lucena-1 or (**D**) FEPS after the free or inhibited C6-NBD-cer efflux. Black squares or inverted triangles respectively indicate Lucena-1 or FEPS untreated cells, and white ones, VP or MK-571 treatment, with lines indicating the median of each population. n=4, with **\*p<0.05** and \***\*p<0.01**.

## Discussion

A number of studies successfully demonstrated associations linking glycosphingolipids and multidrug resistance in several solid tumor types through overexpression of UGCG (23), though few addressed how would its inhibition affect both the expression and efflux activity of ABCB1 and ABCC1 on non-polarized cells. There is evidence that the transport of membrane lipid analogs varies in cellular localization; although ABCB1 translocates C6-NBD-sphingolipids across the apical plasma membrane, ABCC1 seems to transport those analogs to the basolateral plasma membrane on the polarized kidney cell LLC-PK1 (20). Observations that are more recent indicate that ABCB1 and ABCC1 differ in cellular localization, and ABCB1 was shown to transport ceramides along Golgi membranes as well (18). Albeit these observations suggest a similar aptitude to transport sphingolipids, the cellular and organ expressions of ABCB1 and ABCC1 likely take parts on how diverse cells manage sphingolipid levels during proliferation, differentiation, apoptosis and response to cellular stresses. Cells from epithelial origin may express ABC proteins in opposite sides of the cell membrane in a way that, for both blood-brain barrier cells and in the placenta, ABCB1 and ABCC1 are considered, respectively, apical and basolateral transporters (24, 25). Our study proposes, given that peripheral leukemic cells from hematopoietic origins do not present this organization, that ABCB1 and ABCC1 might play complementary yet diverse roles in dealing with xenobiotics and/or sphingolipid mediators.

In a normal bone marrow, it has been demonstrated that GSL levels vary among the distinct stages of erythrocyte differentiation, in a way that GM3 is increased on more differentiated cells such as megakaryocytes (26). Likewise, the efflux activity mediated by ABC transporters is important for the development of hematopoietic progenitors, since ABCB1 and ABCC1 are present on cells with undifferentiated phenotypes on bone marrow or peripheral blood from human (27, 28) and murine (29, 30) origins. Chronic myeloid leukemias result from a reciprocal translocation of BCR and ABL genes among chromosomes 9 and 22 during the erythroblast stage, forming the Philadelphia chromosome. This abnormality contains the chimeric BCR-ABL oncogene, which encodes a protein with constitutive tyrosine kinase activity (31) that sustains proliferative cell signaling and evasion of apoptosis through increased membrane GM1 among other phenotypes (32). Therapy is performed with tyrosine kinase inhibitors such as imatinib mesylate in combination with standard chemotherapeutic drugs for remission (33). Among 30% of patients display some degree of resistance to these drugs, which is closely associated to the expression and activity of ABCB1 (34).

Despite originating from the highly undifferentiated erythroblastic K562 cell line, Lucena-1 and FEPS MDR cells used in this work present higher number of megakaryocytes as well as more differentiated profiles (35). Conversely, it was reported that K562 cells selected for resistance with minimal concentrations of vinblastine and epirubicin, drugs that share similarities in structure and in modes of action with VCR and DNR, showed increased ABCB1 expression with no correlation to specific markers of erythroid origin (36). Considering this, we used different assays to assess GSL and UGCG expression. Our results indicated that selection with the chemotherapeutic drugs VCR or DNR led to a complex and diversified GSL profile, despite UGCG was only significantly increased on FEPS. In this context, treatment with the UGCG inhibitor EtDO-P4 dose-dependently reduced viability on all three cells, nonetheless only Lucena-1 and FEPS showed significant reductions in GM1 levels. A closer observation of the histograms of treated and control cells suggests that MDR cells are more homogenous in GM1 contents than the parental K562. In parallel, both MDR leukemias were sensitized to DNR, Lucena-1 to VCR and FEPS to CDDP as well. This may relate to the specific mechanism of action exerted by each chemotherapeutic during selection, since small differences in expression and in the ceramide portions of sialylated glycolipids could be observed between Lucena-1 and FEPS. Doxorubicin, a DNR analog, was demonstrated to increase ceramide production and UGCG expression on an assortment of tumor cells with variable degrees of drug resistance (37), thus strengthening the relationship between sphingolipids and drug resistance.

Pharmacological UGCG inhibition could be increasing ceramide levels, which along with the stress caused by chemotherapeutics, would explain the higher sensitivity of MDR cells. K562 would possibly shift ceramide glycosylation to galactosylceramide (GalCer) rather than GlcCer, a fact that was described to happen on U-937 and HL-60 human leukemic cells, when the EtDO-P4 analogs PPMP and PDMP actively protected those cells to DNR toxicity (38). In accordance with our results, MDR cells showed lower mitochondrial membrane potentials (Δψ_m_) when GSL depletion was concomitant with DNR treatment, with toxic EtDO-P4 concentrations or when C6-cer was added to the cultures. In those conditions, ceramides could interact with energized mitochondria through direct physical contact or by inducing conformational changes on Bax pro-apoptotic protein leading to intrinsic pathway of apoptosis or caspase-independent cell death (39, 40). Though our results would not discern those mechanisms the participation of mitochondria in the apoptosis induced after EtDO-P4 is clear, considering that the Δψ_m_ loss observed after 24 h would translate into reduced viability, phosphatidylserine exposure and DNA fragmentation after 72 h.

The equilibrium of ceramides and GlcCer, GalCer or GSL may influence cell fate and in extension, the success of therapeutic interventions, and a variety of drug-resistant cell lines present increased cholesterol, sphingomyelin and GSL in comparison to their respective sensitive counterparts (41). The MDR phenotype of cells used in this work is well described, being representative of freshly obtained cells from patients with chemotherapy refractory chronic myeloid leukemia (42, 43). In spite of data indicating that GlcCer modulates ABCB1 expression (17), and considering the promiscuity and overlap in recognition of substrates (44), the relevance of GSL to ABC activity is controversial at best (45). As such, we observed that sub-toxic GSL depletion was not able to affect the expression of ABCB1 and ABCC1 on neither MDR cell; a 24 h incubation with 2 μM EtDO-P4, however, showed reductions in the expression of ABCB1 exclusively. Conversely, the effect of GSL depletion on the efflux performed by ABCB1 or ABCC subfamily members was the opposite; the fluorescence for CF retained inside Lucena-1 and FEPS pre-treated with EtDO-P4 was consistently higher after the efflux assays, indicating that ABCC transport was partially hindered but not the one mediated by ABCB1.

Lucena-1 and FEPS are cross-resistant to a diversity of compounds with natural and synthetic origins owing to, but not limited to, its high efflux activity mediated by the ABC transporters ABCB1 and ABCC1, and to imatinib mesylate as well, which has been demonstrated to increase ceramide and reduce sphingosine-1-phosphate levels on chronic myeloid leukemias (46). Resistance to that inhibitor associates to expression of ABCB1, however with no clear correlation to its efflux activity (47), which suggests that these cells manage their ceramides in alternative forms, involving or not ABCC1 in specific. In this context, ABCB1-mediated efflux activity was demonstrated to be disconnected from the translocation of ceramides on Golgi (48), and the ABCC1 inhibitor MK-571 was shown to affect retrograde membrane transport and to reduce GlcCer formation (49), evidences that point to at least partial involvement of ABCC in regulating sphingolipid levels. Albeit our results concerning ABCC inhibition could not be directly associated to ABCC1 since CFDA, PRB and MK-571 act as substrates and inhibitors to other members from ABCC subfamily (50), to the best of our knowledge, neither Lucena-1 nor FEPS express ABCC2 or express in significantly lower levels than ABCC1 (51). The participation of other ABCC subfamily members such as ABCC3 and ABCC4, however, could not be discarded and should be taken into account in future studies.

It is somewhat difficult to find suitable combinations of substrates and inhibitors to probe the efflux mediated by each subfamily member and, as such, we opted for the most studied and widely employed ones. CFDA, as stressed in our work, may be transported by other ABCC subfamily members than ABCC1, but not by ABCB1. In the same context, a study performed by Dogan *et al*. in 2014, observed the same outcome on 11 leukemia cell lines with diverse degrees of drug resistance (50). That study considered CFDA or calcein the best substrates to prime ABCC1 activity, and given that calcein is a well-known ABCB1 susbtrate (52), our choice of substrates is in accordance with the literature. The same logic applies to the inhibitors. VP, shown to present ABCC1 modulation properties (53), was selective to ABCB1 when combined with Rho123 as substrate and did not affect CFDA efflux. MK-571 on the other hand is a potent, specific inhibitor of ABCC subfamily proteins that has no effect on ABCB1 (50). It is important to notice that we employed PRB as opposed to the more specific MK-571 ABCC inhibitor during the efflux assays on EtDO-P4 pre-treated cells since it could interfere with ceramide levels (49). As such, when combined to the aforementioned substrates, VP (for ABCB1) and PRB or MK-571 (for ABCC) minimize the possibility of cross-interference.

When C6-cer, one possible subproduct from UGCG inhibition and a known mediator of cell stress, was evaluated as a competitor to either ABCB1 or ABCC transport results were similar to the one after EtDO-P4 treatment, provided that this sphingolipid impaired the ABCC-mediated CF efflux from Lucena-1 and FEPS and increased percentages of CF+ cells after the challenge. One more time, significant changes in neither ABCB1-mediated Rho123 efflux nor Rho123+ cells were observed. Finally, when a fluorescent derivative of C6-cer was employed as sole substrate, once more the MFI and the percentages of C6-NBD-cer+ were significantly higher when ABCC activity was inhibited, with a marginal increase in presence of a ABCB1 inhibitor. It is important to disclose that the efflux assays with C6-cer were performed on shorter (1 h) times than viability assays (72 h), and with a higher cell count as well (2×10^5^ cells on the first; 2×10^4^ cells.mL^-1^ on the latter).

Noteworthy, the effects of C6-cer and EtDO-P4 on MDR cells seemed inversely correlated to K562. Lucena-1 and FEPS showed greater decreases in GM1 contents, moderate resistance to EtDO-P4 cytotoxicity and sensitivity to C6-cer-induced viability loss. Although sharing a common precursor, the chemotherapy employed during selection resulted in distinctive phenotypes, in a way that a microarray analysis showed 130 genes with altered expression comparing K562 and Lucena-1, 932 between K562 and FEPS, and 1211 between the two MDR lines (51), ABCB1 being the most overexpressed gene in these cells. Considering this, it was beyond our focus to thoroughly investigate the specific way each cell line would respond to UGCG inhibition; nonetheless, our results point to the possibility of exogenous C6-cer as well as the combination of EtDO-P4 and standard drugs inducing collateral sensitivity, a hypersensitivity towards secondary agents that arises from the development of resistance towards an unrelated primary drug. Those agents passively enter the cell and, after being extruded by ABCB1 or ABCC1, repeat this futile cycle, increasing ATP consumption. Replenishment of ATP increases oxidative stress and the demand for glutathione, a prime ABCC1 substrate (44) and the main peptide involved on defense to oxidative stress. This ultimately leads cells to a form of synthetic lethality due to depletion of these molecules (54, 55). Downregulation of ABCB1 after UGCG impairment would increase intracellular levels of chemotherapeutics and, as consequence, *de novo* production of ceramides in the ER. In parallel, ABCC1 would mediate the efflux of those sphingolipids and drugs as well, further depleting cells of glutathione and ATP, resulting in Δψ_m_ loss and apoptosis through collateral sensitivity. This possibility is feasible once we consider that treatment with C2-ceramide reduced levels of this peptide on a model of epidermal tumor (56) and induced Δψ_m_ loss and apoptosis on models of prostate and colon cancer (40).

In a broader scope, high levels of plasma circulating ceramides can result from a diversity of inflammatory-associated conditions such as diabetes, obesity and cardiovascular diseases, possibly due to changes in remodeling of membrane sphingolipids (57). Concerning cancer, the first study to demonstrate clinical relevance of ceramides to breast cancer, reported higher ceramide levels in the neoplastic tissue than in peri-tumor region or in the plasma of primary breast cancer patients, associated with better prognostic (58). Strikingly, a few studies related increased long-chain ceramide levels on the plasma from patients with advanced ovarian (59) and pancreatic cancer (60), related to higher malignancy. Therefore, participation of ABC transporters on ceramide homeostasis and in an efflux mechanism to extracellular plasma acceptors such as apolipoproteins, apart from ceramide glycosylation or conversion to sphingomyelin had already been proposed but not demonstrated (61). Given the ubiquitous expression of ABCC1, its active efflux of inflammatory lipid mediators such as leukotriene C4 and sphingosine-1-phospate and its association to poor prognostic in cancer (19, 44), ABCC1 emerges as candidate for this role. All things considered, our data indicate that ABCB1 and ABCC subfamily transporters are differentially downregulated following UGCG inhibition.

In this work, our results extend the relationship between sphingolipid glycosylation and acquired resistance mechanisms on models of human MDR neoplastic cells. Further studies addressing the modulation of sphingolipid levels and their cellular fates could translate in the development of strategies directed to MDR hematologic neoplasias, or to the proposal of an adjuvant, off-label use of drugs currently employed against Gaucher’s disease, which accumulates GlcCer in multiple organs as in the bone marrow. Comprehension of the distinct roles of ABC proteins on the clearance of ceramides could ultimately return therapeutic possibilities to patients with chemotherapy refractory leukemias.

### Experimental procedures

#### Cell lines

The chronic myeloid leukemia cell lines K562, Lucena-1 (K562/VCR) and FEPS (K562/DNR) were cultured in RPMI-1640 medium (Sigma-Aldrich, St Louis, MO, USA) supplemented with 25 mM HEPES and 2 g.L^-1^ sodium bicarbonate adjusted to pH 7.4, 100 U penicillin and 100 μg.mL^-1^ streptomycin (all obtained from Sigma-Aldrich) and with 10% fetal bovine serum (FBS) (Thermo Fischer Scientific, Waltham, MA, USA) inactivated at 56 °C for 1 h prior to use. Dr. Vivian M. Rumjanek kindly donated the MDR cells Lucena-1 and FEPS. Briefly, K562 cells were exposed to increasing concentrations of the chemotherapeutic drugs vincristine sulfate (VCR) and daunorubicin hydrochloride (DNR) (both from Sigma-Aldrich), as described before (62, 63). For subcultures, 2×10^4^ cells.mL^-1^ were harvested every 3 days, complete RPMI was added and then maintained at 37 °C in 5% CO_2_. Lucena-1 and FEPS were cultured, respectively, in the presence of 60 nM VCR and 500 nM DNR in order to maintain the MDR phenotypes. Prior to all experiments, the MDR Lucena-1 and FEPS were cultured free of drugs to avoid additive effects.

#### Extraction, purification and analysis of glycosphingolipids

Total GSL were obtained by previously described procedures (64, 65). K562, Lucena-1 and FEPS (2×10^4^ cells.mL^-1^) were seeded in complete RPMI for 72 h in 150 mm Petri plates in the presence of a sub-toxic concentration of the UGCG inhibitor d-threo-1-(3,4,-ethylenedioxy)phenyl-2-palmitoylamino-3-pyrrolidino-1-propanol (EtDO-P4) (66) (Glixx Laboratories, Hopkinton, MA, USA). Controls were treated with the diluent, 0.1% absolute ethanol (Sigma-Aldrich Brazil, Duque de Caxias, RJ, Brazil). Equal amounts of 5×10^7^ cells were harvested and washed in PBS, and precipitates were extracted with 1 mL isopropanol/hexane/water 55:25:20 (vol/vol/vol, lower phase) twice. The extracts were dried under N_2_ stream and then saponified in 2 mL 0.1 M NaOH/methanol to degrade glycerophospholipids. After extraction with hexane, total GSL were isolated using C_18_ cartridges (Analytichem International/Varian, Harbor City, CA, USA) as described, and analyzed by thin-layer chromatography (TLC) followed by staining with a solution of 6% resorcinol and 1% CuSO_4_ in concentrated HCl for the analysis of sialylated GSL. A commercial GSL standard containing a mix of GM1, GM2 and GM3 was chromatographed as reference (Matreya LLC, State College, PA, USA). All solutions were prepared with HPLC-grade reagents from Sigma-Aldrich, Tedia (Fairfield, OH, USA) or SK Chemicals (Seongnam-si, Gyeonggi-do, South Korea).

#### Western blot analysis

K562 and their MDR counterparts Lucena-1 and FEPS were cultured as described before for 48 h, then lysed in lysis buffer (1% Triton X-100; 150 mM NaCl; 25 mM Tris, pH 7.4; 5 mM EDTA; 0.5% sodium deoxycholate; 0.1% SDS; 5 mM tetrasodium pyrophosphate; 50 mM sodium fluoride; 1 mM sodium orthovanadate) containing 1:200 protease inhibitor cocktail (Sigma-Aldrich), and centrifuged at 21000×g for 10 min at 4 °C. 60 μg protein was subjected to SDS–PAGE, followed by transfer to PVDF membranes (Millipore, Burlington, MA, USA). After blocking with 5% skim milk for 30 min, membranes were rinsed with T-TBS and stained with anti-UGCG (clone 1E5; Santa Cruz Biotech, Dallas, TX, USA) and anti-β-actin (clone AC-74, Sigma-Aldrich) antibodies overnight at 4 °C. Membranes were rinsed, incubated with anti-mouse IgG-HRP (Cell Signaling Technology, Danvers, MA, USA) for 2 h at room temperature. After rinsing with T-TBS, membranes were developed by use of Western Lightning Chemiluminescence Reagent Plus (Perkin Elmer, Waltham, MA, USA). Densitometry analyses were performed using ImageJ 1.52d software (U.S. National Institutes of Health, Bethesda, MA, USA) (67). The relative expression of UGCG was calculated as the density of the UGCG band divided by the density of the β-actin band for each cell line.

#### Assessment of GM1 expression on plasma membranes

The glycosphingolipid monosialotetrahexosylganglioside (GM1) was assessed on cell surfaces by flow cytometry. K562, Lucena-1 and FEPS cells were cultured for 24 h as described before. Cell density was adjusted to 2×10^5^ per well and then incubated with 5 μg.mL^-1^ fluorescein isothiocyanate-conjugated cholera toxin (CHT-FITC, Sigma-Aldrich) for 30 min at 4 °C in a light-protected environment. Cell suspensions were centrifuged, resuspended in cold PBS and then analyzed by flow cytometry. Five identical polypeptide B subunits from CHT specifically bind to the GM1 on cell surfaces, and considering that the GSL biosynthesis occurs in a stepwise fashion, the GM1 MFI can relate to the global cell GSL composition (66).

#### In vitro cell viability

Cell viability was determined with the tetrazolium salt 3-(4,5-dimethylthiazol-2-yl)-2,5-diphenyltetrazolium (MTT; Sigma-Aldrich). Briefly, 2×10^4^ cells.mL^-1^ were treated with a range of concentrations of either EtDO-P4, VCR, DNR, cisplatin (CDDP, Incel; Darrow Laboratórios S/A, Rio de Janeiro, RJ, Brazil) or N-hexanoyl-D-erythro-sphingosine (C6-ceramide, C6-cer; Avanti Polar Lipids, Alabaster, AL, USA) for 72 h at 37 °C in 5% CO_2_. Alternatively, cells were co-incubated with 1 μM EtDO-P4 and a range of concentrations of VCR, DNR or CDDP. Negative controls were prepared with the respective diluents (0.1% absolute ethanol for EtDO-P4 or C6-cer; 0.1% DMSO for VCR and DNR; RPMI for CDDP). Then, MTT was added to each well to a final concentration of 0.5 mg.mL^-1^, and plates were incubated at 37 °C in 5% CO_2_ for 3 h in a light-protected environment. After centrifugation, 200 μL DMSO was added to dissolve the dark-blue formazan crystals formed after MTT reduction. Absorbance was measured on a Beckman Coulter AD340S spectrophotometer microplate reader (Beckman Coulter, Brea, CA, USA) at 570 nm. The concentrations giving half-maximal inhibition (IC_50_) were calculated by non-linear regression using the GraphPad Prism version 7.0 software (GraphPad Software, San Diego, CA, USA).

#### Apoptosis assay

The Annexin V/propidium iodide (PI) assay was performed for apoptosis detection. Lucena-1 and FEPS cells were incubated in 24-well plates in the conditions described above and treated with EtDO-P4. After 72 h, cell density was adjusted to 5×10^5^ cells per sample, washed with phosphate-buffered saline (PBS) supplemented with 5% FBS and resuspended in a solution of Annexin V-FITC and propidium iodide (PI) (BD Biosciences, San Diego, CA, USA) in accordance with the manufacturer’s protocol. Cells were incubated at room temperature for 15 min and analyzed by flow cytometry. Dot-plots were divided into four quadrants as follows: upper left (PI+/Annexin-V-), necrotic cells; upper right (PI+/Annexin-V+), late apoptotic cells; lower left (PI-/Annexin-V-), viable cells; lower right (PI-/Annexin-V+), early apoptotic cells.

#### Assessment of mitochondrial membrane potential

Changes in the mitochondrial membrane potential (Δψ_m_) were probed using the rhodamine 123 dye (Rho123; Sigma-Aldrich), which specifically stains energized mitochondria in cultured cells (68). Lucena-1 and FEPS cells were cultured for 24 h with DNR, EtDO-P4 or C6-cer as described before. Following, cell density was adjusted to 2×10^5^ per sample and then incubated with 2.5 μM Rho123 for 30 min at 37 °C in 5% CO_2_ in a light-protected environment. Cell suspensions were centrifuged, resuspended in cold PBS and then analyzed by flow cytometry.

#### Immunophenotypes of MDR cells

The expression of the ABC transporters ABCB1 and ABCC1 was evaluated by flow cytometry. Lucena-1 and FEPS were treated with EtDO-P4 for 24 h as described before, cell density was adjusted to 2×10^5^ per sample at room temperature, then cells were permeabilized and fixed with BD FACS Lysing solution (BD Biosciences) for 10 min and non-specific binding was blocked with PBS 10% FBS for 20 min. Following incubation with anti-ABCB1 (clone D-11) or anti-ABCC1 (clone QCRL-1) human primary antibodies (both from Santa Cruz Biotech, Dallas, Texas, USA) for 30 min at 4 °C, cells were washed with cold PBS and stained with Alexa488-conjugated mouse anti-human IgG secondary antibody (Thermo Fischer Scientific) for 30 min at 4 °C in a light-protected environment. Cell suspensions were centrifuged, resuspended in cold PBS and then analyzed on a flow cytometer.

#### ABC-mediated efflux assays

The ABCB1 and ABCC transport assays were performed, respectively, with the use of the Rho123 and 5(6)-carboxyfluorescein diacetate (CFDA) dyes (Sigma-Aldrich). Rho123 and CFDA are able to passively distribute into the cell, and while the first is fluorescent and is actively extruded by ABCB1, the latter undergoes hydrolysis by nonspecific esterases in the cytosol, originates the fluorescent substrate carboxyfluorescein (CF) that only then is transported out by ABCC subfamily members, notably ABCC1 (69). Briefly, assays were performed in two 30-minute steps, sufficient for the accumulation and efflux of dyes, carried on at 37 °C in 5% CO_2_ in a light-protected environment. 2×10^4^ cells.mL^-1^ were treated for 24 h with 1 μM EtDO-P4 as described prior to the assays. Then, 2×10^5^ Lucena-1 or FEPS cells were incubated in 96-well plates with 250 nM Rho 123 or 500 nM CFDA diluted in RPMI medium to allow accumulation of the dyes within cells. Following, cells were centrifuged at 200×g for 7 min and resuspended in fresh RPMI medium to allow efflux of the dyes (free efflux). In parallel, cells were incubated with the ABCB1 inhibitor verapamil (VP) (70) or the ABCC inhibitors probenecid (PRB) or MK-571 (50) (inhibited efflux), respectively at the concentrations of 10 μM, 1.25 mM and 25 μM. As negative control, cells were exposed to medium only. Next, cells were again centrifuged, resuspended in cold PBS and maintained on ice until acquisition by flow cytometry. Alternatively, C6-cer was employed as a competitive inhibitor during both steps of the assay. Fluorescence histograms were divided into two distinct areas: Rho123 or CF-negative on the left, accounting for 95% of control cells with low MFI for Rho123 or CF, and Rho123 (Rho123+) or CF-positive (CF+) on the right, corresponding to cells still loaded with dyes after efflux phase. A vertical line on each graph indicated those regions.

#### C6-ceramide efflux assays

Assays were performed in two 30-minute steps, in similar conditions to the ABC-mediated efflux assays. 2×10^5^ Lucena-1 or FEPS cells were incubated in 96-well plates with 1 μM nitrobenzoxadiazole-labeled C6-ceramide (C6-NBD-cer) (Avanti Polar Lipids) diluted in 0.1% absolute ethanol to allow accumulation of the fluorescent sphingolipid within cells. In parallel, cells were incubated with the ABCB1 inhibitor verapamil (VP) or the ABCC inhibitor MK-571 (inhibited efflux). As negative control, cells were exposed to medium only, and treated as described before for acquisition by flow cytometry. Fluorescence histograms were divided into two distinct areas: C6-NBD-cer-negative on the left, accounting for 95% of control cells with low MFI, and C6-NBD-cer-positive (C6-NBD-cer+) on the right, corresponding to cells still loaded with the fluorescent sphingolipid after efflux phase. A vertical line on each graph indicated those regions.

#### Flow cytometry

The median fluorescence intensities (MFI) from 15,000 viable cells, gated in accordance with forward (FSC) and side scatter (SSC) parameters representative of cell size and granularity, were acquired using the FL1-H filter on a BD FACSCalibur flow cytometer (BD Biosciences). All post-analyses were performed on Summit version 4.3 software (Dako Colorado, Inc., Fort Collins, CO, USA).

#### Statistical analysis

Statistical analyses were performed using GraphPad Prism version 7.0 software. For two paired comparisons, statistical significance was calculated by parametric Student’s t-tests for normally distributed data, according to the D’Agostino-Pearson test. Otherwise, the non-parametric Wilcoxon test was employed. For more than two comparisons, unpaired one-way ANOVA or Kruskal-Wallis tests were used, respectively, for parametric and nonparametric data. Otherwise, paired one-way ANOVA and the Friedman test were employed for parametric and nonparametric data respectively. Bonferroni’s post-test was used for parametric data while the Dunn’s post-test was employed for nonparametric data. Null hypotheses were rejected when p-values were lower than 0.05, and significances were represented by (**\***) for p <0.05, (**\*\***) for p <0.01 and (**\*\*\***) for p <0.001.

## Acknowledgements

The authors would like to thank Prof. Vivian M. Rumjanek for providing Lucena-1 and FEPS cells and Guilherme G. Fonseca for technical assistance during part of this work.

## FOOTNOTES

Funding was provided by the Conselho Nacional de Desenvolvimento Científico e Tecnológico – CNPq; Financiadora de Estudos e Projetos – FINEP; Fundação do Câncer - Programa de Oncobiologia; Fundação Carlos Chagas Filho de Amparo à Pesquisa do Estado do Rio de Janeiro – FAPERJ; Coordenação de Aperfeiçoamento de Pessoal de Nível Superior – CAPES.

The abbreviations used are: MDR, multidrug resistance; GlcCer, glucosylceramide; GSL, glycosphingolipid; UGCG, UDP-glucose ceramide glucosyltransferase; ABCB1/Pgp, P-glycoprotein; ABCC1/MRP1, multidrug resistance-associated protein 1; GSH, glutathione; EtDO-P4, D-threo-1-(3,4,-ethylenedioxy)phenyl-2-palmitoylamino-3-pyrrolidino-1-propanol; GM1, monosialotetrahexosylganglioside; CHT-FITC, fluorescein isothiocyanate-conjugated cholera toxin; VCR, vincristine; DNR, daunorubicin; CDDP, cisplatin; C6-cer, N-hexanoyl-D-erythro-sphingosine; Δψ_m_, mitochondrial membrane potential; CCCP, Carbonyl cyanide 3-chlorophenylhydrazone; PI, propidium iodide; Rho123, rhodamine 123; CFDA, 5(6)-carboxyfluorescein diacetate; CF, carboxyfluorescein; VP, verapamil; PRB, probenecid; MK-571, 5-(3-(2-(7-Chloroquinolin-2-yl)ethenyl)phenyl)-8-dimethylcarbamyl-4,6-dithiaoctanoic acid sodium salt hydrate; C6-NBD-cer, N-[6-[(7-nitro-2-1,3-benzoxadiazol-4-yl)amino]hexanoyl]-D-erythro-sphingosine; MFI, median fluorescence intensity; GalCer, galactosylceramide.

## REFERENCES

1. Wu, C. P., Hsieh, C. H., and Wu, Y. S. (2011) The emergence of drug transporter-mediated multidrug resistance to cancer chemotherapy. Mol Pharm 8, 1996–2011

2. Ween, M. P., Armstrong, M. A., Oehler, M. K., and Ricciardelli, C. (2015) The role of ABC transporters in ovarian cancer progression and chemoresistance. Crit. Rev. Oncol. Hematol. 96, 220–256

3. Hummel, I., Klappe, K., Ercan, C., and Kok, J. W. (2011) Multidrug resistance-related protein 1 (MRP1) function and localization depend on cortical actin. Mol. Pharmacol. 79, 229–240

4. Meszaros, P., Hummel, I., Klappe, K., Draghiciu, O., Hoekstra, D., and Kok, J. W. (2013) The function of the ATP-binding cassette (ABC) transporter ABCB1 is not susceptible to actin disruption. Biochim. Biophys. Acta 1828, 340–351

5. Leslie, E. M., Deeley, R. G., and Cole, S. P. (2001) Toxicological relevance of the multidrug resistance protein 1, MRP1 (ABCC1) and related transporters. Toxicology 167, 3–23

6. Sharom, F. J. (2014) Complex Interplay between the P-Glycoprotein Multidrug Efflux Pump and the Membrane: Its Role in Modulating Protein Function. Front Oncol 4, 41

7. Cole, S. P., Bhardwaj, G., Gerlach, J. H., Mackie, J. E., Grant, C. E., Almquist, K. C., Stewart, A. J., Kurz, E. U., Duncan, A. M., and Deeley, R. G. (1992) Overexpression of a transporter gene in a multidrug-resistant human lung cancer cell line. Science 258, 1650–1654

8. Dyatlovitskaya, E. V., and Kandyba, A. G. (2006) Role of biologically active sphingolipids in tumor growth. Biochemistry (Mosc). 71, 10–17

9. Saddoughi, S. A., and Ogretmen, B. (2013) Diverse functions of ceramide in cancer cell death and proliferation. Adv. Cancer Res. 117, 37–58

10. Senchenkov, A., Litvak, D. A., and Cabot, M. C. (2001) Targeting ceramide metabolism--a strategy for overcoming drug resistance. J. Natl. Cancer Inst. 93, 347–357

11. Stefanovic, M., Tutusaus, A., Martinez-Nieto, G. A., Barcena, C., de Gregorio, E., Moutinho, C., Barbero-Camps, E., Villanueva, A., Colell, A., Mari, M., Garcia-Ruiz, C., Fernandez-Checa, J. C., and Morales, A. (2016) Targeting glucosylceramide synthase upregulation reverts sorafenib resistance in experimental hepatocellular carcinoma. Oncotarget 7, 8253–8267

12. Gencer, E. B., Ural, A. U., Avcu, F., and Baran, Y. (2011) A novel mechanism of dasatinib-induced apoptosis in chronic myeloid leukemia; ceramide synthase and ceramide clearance genes. Ann. Hematol. 90, 1265–1275

13. Zhao, S., Yang, Y. N., and Song, J. G. (2004) Ceramide induces caspase-dependent and - independent apoptosis in A-431 cells. J. Cell. Physiol. 199, 47–56

14. Hannun, Y. A., and Obeid, L. M. (2008) Principles of bioactive lipid signalling: lessons from sphingolipids. Nat Rev Mol Cell Biol 9, 139–150

15. Kartal Yandim, M., Apohan, E., and Baran, Y. (2013) Therapeutic potential of targeting ceramide/glucosylceramide pathway in cancer. Cancer Chemother. Pharmacol. 71, 13–20

16. Liu, Y., Xie, K. M., Yang, G. Q., Bai, X. M., Shi, Y. P., Mu, H. J., Qiao, W. Z., Zhang, B., and Xie, P. (2010) GCS induces multidrug resistance by regulating apoptosis-related genes in K562/AO2 cell line. Cancer Chemother. Pharmacol. 66, 433–439

17. Liu, Y. Y., Gupta, V., Patwardhan, G. A., Bhinge, K., Zhao, Y., Bao, J., Mehendale, H., Cabot, M. C., Li, Y. T., and Jazwinski, S. M. (2010) Glucosylceramide synthase upregulates MDR1 expression in the regulation of cancer drug resistance through cSrc and beta-catenin signaling. Mol Cancer 9, 145

18. Eckford, P. D., and Sharom, F. J. (2005) The reconstituted P-glycoprotein multidrug transporter is a flippase for glucosylceramide and other simple glycosphingolipids. Biochem. J. 389, 517–526

19. Yamada, A., Nagahashi, M., Aoyagi, T., Huang, W. C., Lima, S., Hait, N. C., Maiti, A., Kida, K., Terracina, K. P., Miyazaki, H., Ishikawa, T., Endo, I., Waters, M. R., Qi, Q., Yan, L., Milstien, S., Spiegel, S., and Takabe, K. (2018) ABCC1-Exported Sphingosine-1-phosphate, Produced by Sphingosine Kinase 1, Shortens Survival of Mice and Patients with Breast Cancer. Mol Cancer Res 16, 1059–1070

20. Raggers, R. J., van Helvoort, A., Evers, R., and van Meer, G. (1999) The human multidrug resistance protein MRP1 translocates sphingolipid analogs across the plasma membrane. J. Cell Sci. 112 (Pt 3), 415–422

21. Klappe, K., Hinrichs, J. W., Kroesen, B. J., Sietsma, H., and Kok, J. W. (2004) MRP1 and glucosylceramide are coordinately over expressed and enriched in rafts during multidrug resistance acquisition in colon cancer cells. Int. J. Cancer 110, 511–522

22. Daniotti, J. L., Lardone, R. D., and Vilcaes, A. A. (2015) Dysregulated Expression of Glycolipids in Tumor Cells: From Negative Modulator of Anti-tumor Immunity to Promising Targets for Developing Therapeutic Agents. Front Oncol 5, 300

23. Wegner, M. S., Gruber, L., Mattjus, P., Geisslinger, G., and Grosch, S. (2018) The UDP-glucose ceramide glycosyltransferase (UGCG) and the link to multidrug resistance protein 1 (MDR1). BMC Cancer 18, 153

24. Evseenko, D. A., Paxton, J. W., and Keelan, J. A. (2007) Independent regulation of apical and basolateral drug transporter expression and function in placental trophoblasts by cytokines, steroids, and growth factors. Drug metabolism and disposition: the biological fate of chemicals 35, 595–601

25. Gazzin, S., Strazielle, N., Schmitt, C., Fevre-Montange, M., Ostrow, J. D., Tiribelli, C., and Ghersi-Egea, J. F. (2008) Differential expression of the multidrug resistance-related proteins ABCB1 and ABCC1 between blood-brain interfaces. J. Comp. Neurol. 510, 497–507

26. Nakamura, M., Kirito, K., Yamanoi, J., Wainai, T., Nojiri, H., and Saito, M. (1991) Ganglioside GM3 can induce megakaryocytoid differentiation of human leukemia cell line K562 cells. Cancer Res. 51, 1940–1945

27. Abbaszadegan, M. R., Futscher, B. W., Klimecki, W. T., List, A., and Dalton, W. S. (1994) Analysis of multidrug resistance-associated protein (MRP) messenger RNA in normal and malignant hematopoietic cells. Cancer Res. 54, 4676–4679

28. Drach, D., Zhao, S., Drach, J., Mahadevia, R., Gattringer, C., Huber, H., and Andreeff, M. (1992) Subpopulations of normal peripheral blood and bone marrow cells express a functional multidrug resistant phenotype. Blood 80, 2729–2734

29. Costa, K. M. D., Valente, R. C., Silva, J., Paiva, L. S., and Rumjanek, V. M. (2018) Glucocorticoid susceptibility and in vivo ABCB1 activity differ in murine B cell subsets. An. Acad. Bras. Cienc. 90, 3081–3097

30. Kyle-Cezar, F., Echevarria-Lima, J., dos Santos Goldenberg, R. C., and Rumjanek, V. M. (2007) Expression of c-kit and Sca-1 and their relationship with multidrug resistance protein 1 in mouse bone marrow mononuclear cells. Immunology 121, 122–128

31. Krusch, M., and Salih, H. R. (2011) Effects of BCR-ABL inhibitors on anti-tumor immunity. Curr. Med. Chem. 18, 5174–5184

32. Cebo, C., Da Rocha, S., Wittnebel, S., Turhan, A. G., Abdelali, J., Caillat-Zucman, S., Bourhis, J. H., Chouaib, S., and Caignard, A. (2006) The decreased susceptibility of Bcr/Abl targets to NK cell-mediated lysis in response to imatinib mesylate involves modulation of NKG2D ligands, GM1 expression, and synapse formation. J. Immunol. 176, 864–872

33. Bixby, D., and Talpaz, M. (2011) Seeking the causes and solutions to imatinib-resistance in chronic myeloid leukemia. Leukemia 25, 7–22

34. Maia, R. C., Vasconcelos, F. C., Souza, P. S., and Rumjanek, V. M. (2018) Towards Comprehension of the ABCB1/P-Glycoprotein Role in Chronic Myeloid Leukemia. Molecules 23

35. Carrett-Dias, M., Almeida, L. K., Pereira, J. L., Almeida, D. V., Filgueira, D. M., Marins, L. F., Votto, A. P., and Trindade, G. S. (2016) Cell differentiation and the multiple drug resistance phenotype in human erythroleukemic cells. Leuk. Res. 42, 13–20

36. Marks, D. C., Davey, M. W., Davey, R. A., and Kidman, A. D. (1993) Differentiation and multidrug resistance in response to drug treatment in the K562 human leukaemia cell line. Br. J. Haematol. 84, 83–89

37. Liu, Y. Y., Yu, J. Y., Yin, D., Patwardhan, G. A., Gupta, V., Hirabayashi, Y., Holleran, W. M., Giuliano, A. E., Jazwinski, S. M., Gouaze-Andersson, V., Consoli, D. P., and Cabot, M. C. (2008) A role for ceramide in driving cancer cell resistance to doxorubicin. FASEB J. 22, 2541–2551

38. Grazide, S., Terrisse, A. D., Lerouge, S., Laurent, G., and Jaffrezou, J. P. (2004) Cytoprotective effect of glucosylceramide synthase inhibition against daunorubicin-induced apoptosis in human leukemic cell lines. J. Biol. Chem. 279, 18256–18261

39. Chang, K. T., Anishkin, A., Patwardhan, G. A., Beverly, L. J., Siskind, L. J., and Colombini, M. (2015) Ceramide channels: destabilization by Bcl-xL and role in apoptosis. Biochim. Biophys. Acta 1848, 2374–2384

40. von Haefen, C., Wieder, T., Gillissen, B., Starck, L., Graupner, V., Dorken, B., and Daniel, P. T. (2002) Ceramide induces mitochondrial activation and apoptosis via a Bax-dependent pathway in human carcinoma cells. Oncogene 21, 4009–4019

41. Ramu, A., Glaubiger, D., and Weintraub, H. (1984) Differences in lipid composition of doxorubicin-sensitive and -resistant P388 cells. Cancer Treat. Rep. 68, 637–641

42. Maia, R. C., Vasconcelos, F. C., de Sa Bacelar, T., Salustiano, E. J., da Silva, L. F., Pereira, D. L., Moellman-Coelho, A., Netto, C. D., da Silva, A. J., Rumjanek, V. M., and Costa, P. R. (2011) LQB-118, a pterocarpanquinone structurally related to lapachol [2-hydroxy-3-(3-methyl-2-butenyl)-1,4-naphthoquinone]: a novel class of agent with high apoptotic effect in chronic myeloid leukemia cells. Invest. New Drugs 29, 1143–1155

43. Salustiano, E. J., Netto, C. D., Fernandes, R. F., da Silva, A. J., Bacelar, T. S., Castro, C. P., Buarque, C. D., Maia, R. C., Rumjanek, V. M., and Costa, P. R. (2010) Comparison of the cytotoxic effect of lapachol, alpha-lapachone and pentacyclic 1,4-naphthoquinones on human leukemic cells. Invest. New Drugs 28, 139–144

44. Cole, S. P. (2014) Multidrug resistance protein 1 (MRP1, ABCC1), a “multitasking” ATP-binding cassette (ABC) transporter. J. Biol. Chem. 289, 30880–30888

45. Dijkhuis, A. J., Klappe, K., Kamps, W., Sietsma, H., and Kok, J. W. (2006) Gangliosides do not affect ABC transporter function in human neuroblastoma cells. J. Lipid Res. 47, 1187–1195

46. Baran, Y., Salas, A., Senkal, C. E., Gunduz, U., Bielawski, J., Obeid, L. M., and Ogretmen, B. (2007) Alterations of ceramide/sphingosine 1-phosphate rheostat involved in the regulation of resistance to imatinib-induced apoptosis in K562 human chronic myeloid leukemia cells. J. Biol. Chem. 282, 10922–10934

47. da Cunha Vasconcelos, F., Mauricio Scheiner, M. A., Moellman-Coelho, A., Mencalha, A. L., Renault, I. Z., Rumjanek, V. M., and Maia, R. C. (2016) Low ABCB1 and high OCT1 levels play a favorable role in the molecular response to imatinib in CML patients in the community clinical practice. Leuk. Res. 51, 3–10

48. Pallis, M., and Russell, N. (2000) P-glycoprotein plays a drug-efflux-independent role in augmenting cell survival in acute myeloblastic leukemia and is associated with modulation of a sphingomyelin-ceramide apoptotic pathway. Blood 95, 2897–2904

49. Kok, J. W., Babia, T., Filipeanu, C. M., Nelemans, A., Egea, G., and Hoekstra, D. (1998) PDMP blocks brefeldin A-induced retrograde membrane transport from golgi to ER: evidence for involvement of calcium homeostasis and dissociation from sphingolipid metabolism. J. Cell Biol. 142, 25–38

50. Dogan, A. L., Legrand, O., Faussat, A. M., Perrot, J. Y., and Marie, J. P. (2004) Evaluation and comparison of MRP1 activity with three fluorescent dyes and three modulators in leukemic cell lines. Leuk. Res. 28, 619–622

51. Moreira, M. A., Bagni, C., de Pinho, M. B., Mac-Cormick, T. M., dos Santos Mota, M., Pinto-Silva, F. E., Daflon-Yunes, N., and Rumjanek, V. M. (2014) Changes in gene expression profile in two multidrug resistant cell lines derived from a same drug sensitive cell line. Leuk. Res. 38, 983–987

52. Legrand, O., Simonin, G., Perrot, J. Y., Zittoun, R., and Marie, J. P. (1999) Both Pgp and MRP1 activities using calcein-AM contribute to drug resistance in AML. Adv. Exp. Med. Biol. 457, 161–175

53. Strouse, J. J., Ivnitski-Steele, I., Waller, A., Young, S. M., Perez, D., Evangelisti, A. M., Ursu, O., Bologa, C. G., Carter, M. B., Salas, V. M., Tegos, G., Larson, R. S., Oprea, T. I., Edwards, B. S., and Sklar, L. A. (2013) Fluorescent substrates for flow cytometric evaluation of efflux inhibition in ABCB1, ABCC1, and ABCG2 transporters. Anal. Biochem. 437, 77–87

54. Lorendeau, D., Dury, L., Nasr, R., Boumendjel, A., Teodori, E., Gutschow, M., Falson, P., Di Pietro, A., and Baubichon-Cortay, H. (2017) MRP1-dependent Collateral Sensitivity of Multidrug-resistant Cancer Cells: Identifying Selective Modulators Inducing Cellular Glutathione Depletion. Curr. Med. Chem. 24, 1186–1213

55. Pluchino, K. M., Hall, M. D., Goldsborough, A. S., Callaghan, R., and Gottesman, M. M. (2012) Collateral sensitivity as a strategy against cancer multidrug resistance. Drug Resist Updat 15, 98–105

56. Davis, M. A., Flaws, J. A., Young, M., Collins, K., and Colburn, N. H. (2000) Effect of ceramide on intracellular glutathione determines apoptotic or necrotic cell death of JB6 tumor cells. Toxicol. Sci. 53, 48–55

57. Haus, J. M., Kashyap, S. R., Kasumov, T., Zhang, R., Kelly, K. R., Defronzo, R. A., and Kirwan, J. P. (2009) Plasma ceramides are elevated in obese subjects with type 2 diabetes and correlate with the severity of insulin resistance. Diabetes 58, 337–343

58. Moro, K., Kawaguchi, T., Tsuchida, J., Gabriel, E., Qi, Q., Yan, L., Wakai, T., Takabe, K., and Nagahashi, M. (2018) Ceramide species are elevated in human breast cancer and are associated with less aggressiveness. Oncotarget 9, 19874–19890

59. Knapp, P., Bodnar, L., Blachnio-Zabielska, A., Swiderska, M., and Chabowski, A. (2017) Plasma and ovarian tissue sphingolipids profiling in patients with advanced ovarian cancer. Gynecol. Oncol. 147, 139–144

60. Jiang, Y., DiVittore, N. A., Young, M. M., Jia, Z., Xie, K., Ritty, T. M., Kester, M., and Fox, T. E. (2013) Altered sphingolipid metabolism in patients with metastatic pancreatic cancer. Biomolecules 3, 435–448

61. Hussain, M. M., Jin, W., and Jiang, X. C. (2012) Mechanisms involved in cellular ceramide homeostasis. Nutr Metab (Lond) 9, 71

62. Daflon-Yunes, N., Pinto-Silva, F. E., Vidal, R. S., Novis, B. F., Berguetti, T., Lopes, R. R., Polycarpo, C., and Rumjanek, V. M. (2013) Characterization of a multidrug-resistant chronic myeloid leukemia cell line presenting multiple resistance mechanisms. Mol. Cell. Biochem. 383, 123–135

63. Rumjanek, V. M., Trindade, G. S., Wagner-Souza, K., de-Oliveira, M. C., Marques-Santos, L. F., Maia, R. C., and Capella, M. A. (2001) Multidrug resistance in tumour cells: characterization of the multidrug resistant cell line K562-Lucena 1. An. Acad. Bras. Cienc. 73, 57–69

64. Freire-de-Lima, L., Gelfenbeyn, K., Ding, Y., Mandel, U., Clausen, H., Handa, K., and Hakomori, S. I. (2011) Involvement of O-glycosylation defining oncofetal fibronectin in epithelial-mesenchymal transition process. Proc. Natl. Acad. Sci. U. S. A. 108, 17690–17695

65. Guan, F., Schaffer, L., Handa, K., and Hakomori, S. I. (2010) Functional role of gangliotetraosylceramide in epithelial-to-mesenchymal transition process induced by hypoxia and by TGF-β. FASEB J. 24, 4889–4903

66. Lee, L., Abe, A., and Shayman, J. A. (1999) Improved inhibitors of glucosylceramide synthase. J. Biol. Chem. 274, 14662–14669

67. Ferreira, T., and Rasband, W. S. (2010-2012) ImageJ User Guide — IJ 1.46.

68. Huang, M., Camara, A. K., Stowe, D. F., Qi, F., and Beard, D. A. (2007) Mitochondrial inner membrane electrophysiology assessed by rhodamine-123 transport and fluorescence. Ann. Biomed. Eng. 35, 1276–1285

69. van der Kolk, D. M., de Vries, E. G., Koning, J. A., van den Berg, E., Muller, M., and Vellenga, E. (1998) Activity and expression of the multidrug resistance proteins MRP1 and MRP2 in acute myeloid leukemia cells, tumor cell lines, and normal hematopoietic CD34+ peripheral blood cells. Clin. Cancer Res. 4, 1727–1736

70. Yusa, K., and Tsuruo, T. (1989) Reversal mechanism of multidrug resistance by verapamil: direct binding of verapamil to P-glycoprotein on specific sites and transport of verapamil outward across the plasma membrane of K562/ADM cells. Cancer Res. 49, 5002–5006

